# Scaling in Biological Systems: A Molecular-Ensemble Duality

**DOI:** 10.1101/2025.01.31.635926

**Authors:** Julie A. Ellsworth, Josh E. Baker

## Abstract

We have observed in muscle the statistical mechanics of irreversible chemical thermodynamics, revealing the solution to multiple seemingly unrelated paradoxes in science. Analogous to Boltzmann’s H theorem, we observe that chemical reaction energy landscapes (ensemble entropic wells) irreversibly evolve over time, pulling reversible chemical reactions forward in time. Loschmidt’s paradox assumes that reversible molecular reactions scale up to irreversible changes in an ensemble, and many mathematical constructs have been created to satisfy this assumption (Boltzmann’s H-function, chemical activities, the kinetics theory of gases, molecular mechanisms of biological function, etc.). However, using a simple statistical argument, here we show that the irreversible time evolutions of molecular and ensemble states are described by two different non-scalable entropies, creating a molecular-ensemble duality in any system on any scale. This inverts common understandings of mechanistic agency and the arrow of time and disproves all molecular mechanisms of irreversible ensemble processes.

**Significance Statement:** This statistical analysis inverts common understandings of mechanistic agency, entropy, and the arrow of time; it solves several paradoxes in physics; and it disproves molecular mechanisms of irreversible processes.

## Introduction

Deterministic physics assumes that the universe follows a single trajectory over time that in theory, with enough information, is described by a single Newtonian master equation. Unpredictable interventions, however, create branch points in this trajectory (1, 2). For example, in thermodynamics the branches of a molecular trajectory are its degrees of freedom where at each branch point a path forward is randomly chosen by thermal energy (3). Propagating branches create disorder over time increasing molecular entropy (4–6). A closed system of molecules equilibrates at maximum disorder and entropy, and once equilibrated, all combinations of branches are equally probable, at which point these trajectories are said to be microscopically reversible (they can return to any state) (7, 8).

This raises an important question. How can molecular reactions that maximize disorder and reversibility determine irreversible directional ensemble processes? Applied to gases, this is Loschmidt’s paradox (9), but the same question applies to chemistry, biology, and other disciplines. Early thermodynamicists knew that a system in which a change in entropy could be mechanically reversed was needed to solve this and other of entropy’s paradoxes (10), and muscle provides us with this system (11–13). In muscle, we have directly observed the statistical mechanics of irreversible chemical thermodynamics. Specifically, we observe that, analogous to Boltzmann’s H theorem, the reaction energy landscape (an entropic well) of molecular (myosin) switches induced by actin binding irreversibly changes with time, pulling the reversible binding reaction forward. In other words, irreversible muscle contraction determines chemistry and not the other way around (13, 14).

Biophysicists assert that the increase in ensemble entropy associated with irreversible muscle contraction does not preclude molecular mechanisms of muscle contraction. Indeed, the assumption that molecules and ensembles are scalable is the rationale for many proposed molecular mechanisms of irreversible function such as Boltzmann’s H-function (15), the kinetic theory of gasses (16), the kinetic theory of chemical reactions, chemical activities (17), and all molecular mechanisms of biological systems (18).

Here, we distill our observations down to a simple statistical analysis and show that the time evolution of molecular switches (branching molecular trajectories) cannot account for the time evolution of an ensemble of switches (all possible branches) because the corresponding entropies are not mathematically scalable. In effect, the molecular branches are entangled in all possible combinations of branches, creating a molecular-ensemble duality. This analysis inverts conventional understandings of entropy, scale, mechanistic agency, and the arrow of time. It solves many of entropy’s paradoxes including Loschmidt’s paradox (9), Gibbs’ paradox (19–21), Maxwell’s demon (22), and Schrodinger’s cat (an entropic paradox), and it disproves all molecular mechanisms of irreversible ensemble processes.

### The Chemistry of Muscle Contraction

Molecular motors in muscle are mechanical switches that reversibly exert force in one state but not the other (23–25). How do these microscopically reversible switches generate irreversible muscle contraction (13)? We have observed in muscle the solution to this and other paradoxes in science. The solution can be expressed mathematically and thus requires no knowledge of molecular switches, thermal energy, biology, chemistry, or physics. Here we present a simple analysis of coins stochastically tossed over time, mirroring Monte Carlo simulations of a two-state chemical relaxation.

### The Kinetics of Coins

If we watch individual coins stochastically tossed at a rate, *k*, we observe coins reversibly flip back and forth from heads, H, to tails, T, at a rate of ½*k* in each direction (Fig. 1A). We can observe numbers of coins, *n*_*T*_ and *n*_*H*_, in states T and H at different instants of time and can look for patterns in the observed time course. However, patterns described in terms of observed unpredictable outcomes, *n*_*T*_ and *n*_*H*_, are correlative, not predictive. For example, a conventional chemical analysis that defines d*n*_*T*_(t)/dt (Fig. 1A, legend) as a function of these parameters is observational, not predictive.

**Fig. 1.**
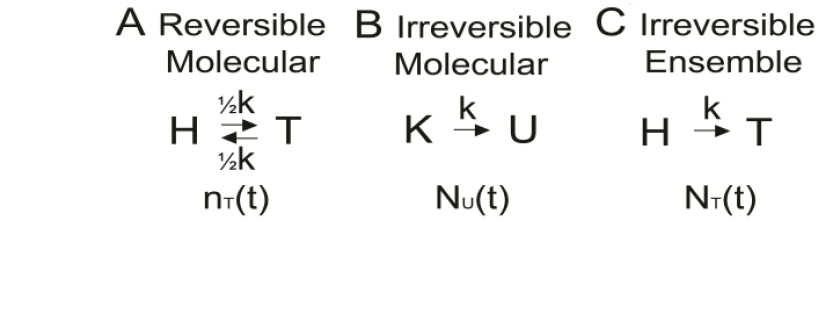
Three kinetic schemes for coins. (A) When individual coins are tossed at a rate, *k*, they reversibly transition between heads, H, and tails, T, at a rate of ½*k* in each direction. The number of coins in state T, *n*_*T*_(t), is observed to reversibly change with time as d*n*_*T*_(t)/dt = – ½*n*_*T*_(t)·*k* + ½*n*_*H*_(t)·*k*. (B) When coins are tossed at a rate, *k*, they irreversibly transition from a known, K, to an unknown, U, state, at a rate, *k*. The number of known coins, *N*_*K*_(t), irreversibly changes with time as d*N*_*K*_/dt = –*N*_*K*_·*k*. (C) When an ensemble of coins is tossed at a rate, *k*, they irreversibly transition from heads, H, to tails, T. The number of coins in state T, *n*_*T*_(t), follows the predictable statistics of the irreversible time course of the microstate, *N*_*T*_(t), which we show changes with time as d(*N*_*H*_ – *N*_*T*_)/dt = (*N*_*H*_ – *N*_*T*_)·*k*.

In a predictive analysis, the time course of tossed coins is irreversible. When a coin in a known state, K, is tossed, it irreversibly transitions at a rate, *k*, to an unknown state, U (Fig. 1B) with two unpredictable branches, H and T. Here, the change with time in the number of known coins, d*N*_*K*_(t)/dt (Fig. 1B, legend) is predictive because we can predict the outcome, U, of a coin toss.

We show below that when we sum up trajectories, *n*_*T*_(t), of multiple coins (an ensemble of coins), the average change with time in the total number, *N*_*T*_(t), of coins in state T follows the same irreversible kinetics as *N*_*U*_(t) (Fig. 1C). For example, a net flux from H to T at a rate, *k*, is observed if *n*_*H*_ > *n*_*T*_. How does this irreversible directional flux from H to T (Fig. 1C) scale up from reversible H to T transitions (Fig. 1A)? The answer is that it does not. We show below that, unlike *n*_*T*_(t), *N*_*T*_(t) is not an actual number of coins. Analogous to Boltzmann’s H theorem, *N*_*T*_(t) is a statistical microstate, or expectation, that describes the evolution with time of the number of possible combinations of branches, H and T, that can result in state *N*_*T*_(t). The irreversible time evolution of the expectation, *N*_*T*_(t), predicts the average *n*_*T*_(t) outcome of stochastically tossed coins. In other words, irreversible ensemble kinetics (Fig. 1C) determines molecular kinetics (Fig. 1A), inverting the agency of molecular determinism. This is the agency we observe in muscle (12, 13) where the irreversible kinetics of an ensemble of molecular switches (i.e., muscle contraction) determines the kinetics of reversible molecular switches.

Unknown complex interactions among molecules are often assumed to be the reason molecular kinetics do not scale up to ensemble kinetics (Loschmidt’s paradox), and molecular mechanisms like Boltzmann’s H-function and chemical activities were created to account for these mysterious interactions. However, the coins above do not interact. In fact, we show below that the reason molecular and ensemble kinetics do not physically scale is because they do not mathematically scale. They are defined by two different entropies (the number of unknown branches, *N*_*U*_, versus the number, *N*_*T*_(t), of possible combinations of branches) that cannot be defined at the same time, creating a molecular-ensemble duality.

There are two formal differences between the kinetics of coin tosses and the kinetics of chemical reactions. First, the simulated stochastic tossing of coins does not require thermal energy, k_B_T. Second, the enthalpic difference between H and T is zero (a non-zero enthalpy changes the probability of H versus T). The statistical analysis below applies equally to the irreversible kinetics of both reactions.

### Molecular and Ensemble Kinetics

We simulated *N* = 10 fair coins stochastically tossed at a rate, *k*, of 1 sec^−1^ all starting as heads. Figure 2A shows these simulations where coin tosses are indicated by red circles and transitions between H (lower) and T (upper) follow the black lines.

**Fig. 2.**
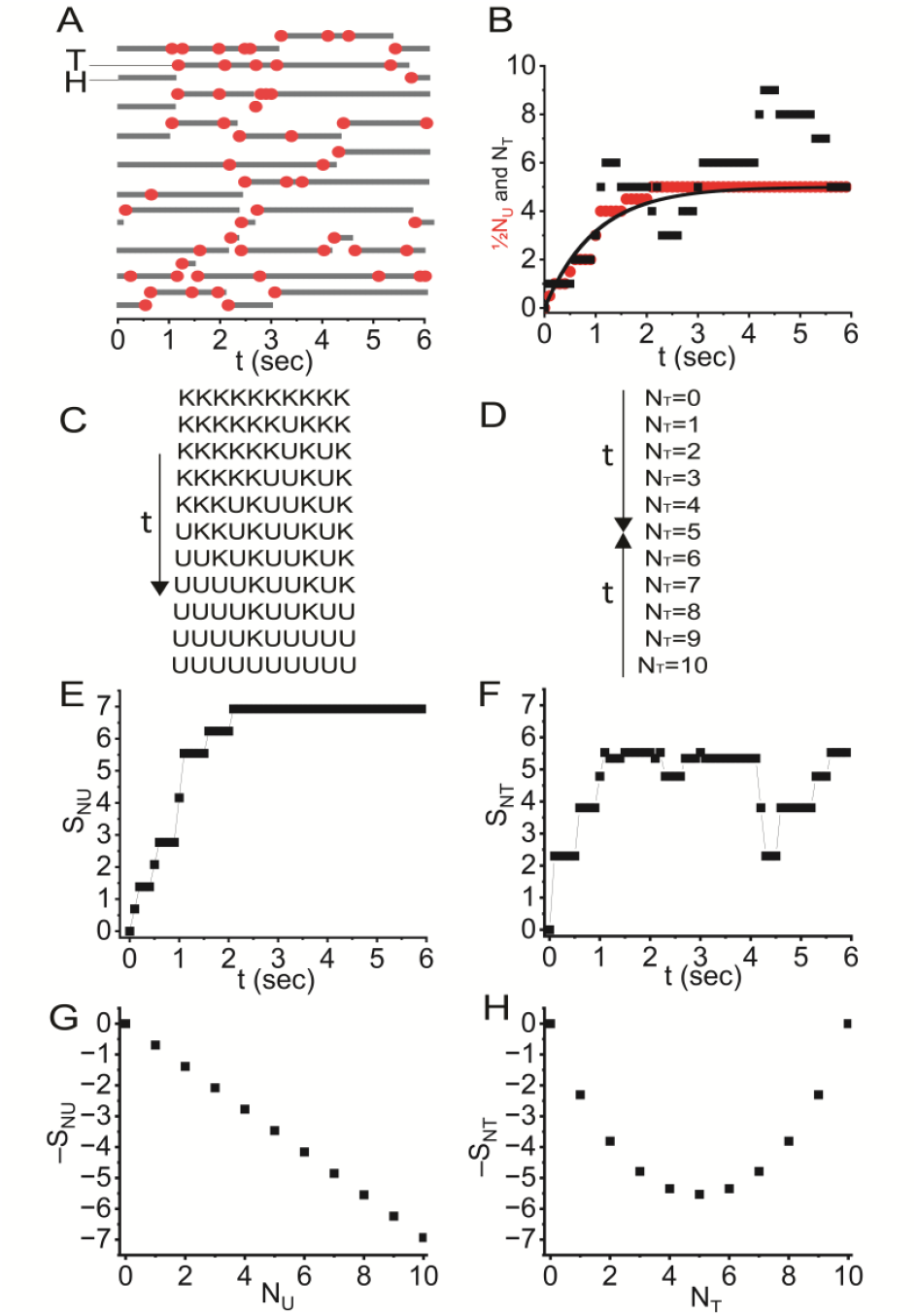
Molecular and ensemble kinetics and entropy. (A) The stochastic tossing of ten coins at a rate, *k*, are simulated (Monte Carlo) with a random number that gives each coin a 10% chance of flipping every 0.1 seconds. A second random number gives a flipped coin a 50% chance of being either heads, H, or tails, T. Transitions between H (lower) and T (upper) are shown (black lines) over time along with coin tosses indicated by red circles. (B) An increase in *N*_*T*_ by ½ at each time-to-first-flip (the first red circle in each trajectory in panel A) is plotted (red circles) along with the sum of all ten trajectories (black squares). The line is *N*_*H*_ = ½*N* – ½*N*·exp(-*k*t). (C) Sequential changes in molecular states from (A) are illustrated. (D) The ensemble states are pulled toward *N*_*T*_ = 5 with time. (E) Molecular entropy, *N*_*u*_·ln(2), calculated from the molecular transient in (B) is plotted. (F) Ensemble entropy, ln[*N*!/(*N*_*H*_!*N*_*T*_!)], calculated from the ensemble transient in (B) is plotted. (G) The *N*_*U*_-dependence of molecular entropy, *N*_*u*_·ln(2), is plotted. (H) The *N*_*T*_-dependence of ensemble entropy, ln[*N*!/(*N*_*H*_!*N*_*T*_!)], is plotted.

An ensemble transient (Fig. 2B, black squares) is obtained by summing all ten trajectories. This transient shows an increase in the number of tails with time, *N*_*T*_(t). An increase in the number, *N*_*U*_(t), of tossed (unknown, U) coins with time (Fig. 2B, red circles) is obtained from the first red circle of each trajectory in Fig. 2A, where *N*_*T*_(t) = ½*N*_*U*_(t) is plotted considering that half of the unknown coins are on average tails. The former describes a flux from H to T (Fig. 1C) while the latter describes the disordering of coins (Fig. 1B). Both transients follow the same irreversible kinetics (Fig. 2B, solid curve). The number of coins, *N*_*U*_, in state U increases with time (Fig. 1B, legend) as *N*_*U*_(t) = *N* – *N*·exp(-*k*t), or *N*_*T*_(t) = ½*N* – ½*N*·exp(-*k*t) (Fig. 2B, solid curve).

While the kinetics, *n*_*T*_(t), of reversible H to T transitions (Fig. 1A) are of interest in chemical analyses, they are unrelated to *N*_*T*_(t) and *N*_*U*_(t). Reversible H to T transitions are averaged away as noise in defining the average ½T in state U. Likewise, *n*_*T*_(t) is the noise in the irreversible ensemble transient, *N*_*T*_(t) (Fig. 2B, black squares). Distilled down to a simple statistical truth, Loschmidt’s paradox states that noise, *n*_*T*_(t), does not determine signal, *N*_*T*_(t). That is, reversible molecular transitions (Fig. 1A) do not determine irreversible ensemble processes (Fig. 1C). This is obviously not a statistical paradox, and it is a scientific paradox only because it is assumed that *n*_*T*_(t) (Fig. 1A) determines *N*_*T*_(t) (Fig. 1C). Here we show that the signal, *N*_*T*_(t) is a statistical microstate in a probability distribution that evolves irreversibly with time, and that the predictable statistics of the time course of *N*_*T*_(t) determines the observed *n*_*T*_(t), not the other way around.

There is one possible way, *W*_*NT=0*_ = 1, for coins to be in state *N*_*T*_ = 0, and ten possible ways, *W*_*NT=1*_ = 10, for coins to be in state *N*_*T*_ = 1. This increase in *W* pulls coins from *N*_*T*_ = 0 to *N*_*T*_ = 1, resulting in a net flux from H to T. Because it can never exceed *N*_*U*_(t), *N*_*T*_(t) is governed by the irreversible time course of *N*_*U*_(t). As *N*_*T*_(t) increases with time, *W*_*NT*_(t) increases with time as

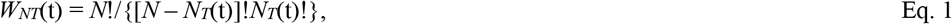

pulling the distribution of coins from H to T with an increase in *N*_*T*_(t). Here we describe the irreversible kinetics of *N*_*T*_(t) that determines the observed kinetics of *n*_*T*_(t) described by conventional chemistry.

### Molecular and Ensemble Entropy

Shannon’s entropy, S, is defined (5) by the number of possible ways, *W*, a state can exist in a closed system:

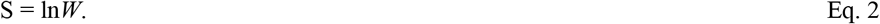

If all ten coins are randomly flipped, then *W*_*NU=0*_ = 2^10^, and S_*NU=10*_ = 10·ln2. However, only a fraction of the maximum entropy and corresponding maximum disorder is contained within a specific state at a given instant of time.

The states *N*_*U*_ and *N*_*T*_ represent two different ways of describing the same system of coins, which means they are associated with two different definitions of entropy. The former describes irreversible transitions down unknown branched pathways (noise) on a scale we refer to as the obscurus (the unknown). The latter describes irreversible changes in the statistics of all possible branches (signal) on a scale we refer to as the omnia possibilia (the all possible).

On the scale of the obscurus, at time t = 0 there is one way (*W*_*NU=0*_ = 1) that all 10 coins are heads. Here, the obscurus entropy, *S*_*NU=0*_, equals zero. After the first K to U transition there are two unpredictable branches (*W*_*NU=1*_ = 2), and *S*_*NU=1*_ = ln2. After the second K to U transition, *S*_*NU=2*_ = 2·ln2. After the *n*^th^ transition, *S*_*NU*_ = *n*·ln2 = *N*_*U*_·ln2 where ln2 is often referred to as one bit of information. That is, one bit of information is lost with each transition from K to U, and after all coins have flipped at least once, the coins equilibrate in a state of maximum disorder and entropy, S_*NU=10*_ = 10·ln2. Figure 2E is a plot of the time course of the obscurus entropy corresponding to the molecular trajectory in Fig. 2B (red circles).

The omnia entropy is a form of Boltzmann’s entropy, S_*NT*_ = ln[*N*!/(*N*_*H*_!*N*_*T*_!)] (Eqs. 1 and 2), only here thermal energy is not specified as the energy that tosses the coins. In Fig. 2F, S_*NT*_ is plotted for the ensemble trajectory in Fig. 2B. Unlike an increase in obscurus entropy, an increase in omnia entropy is not associated with the loss of bits of information; rather, it is associated with an irreversible change in the ensemble state *N*_*T*_(t). We have observed that an increase in the (omnia) entropy of a binary system (Fig. 2F) is the mechanism of irreversible muscle contraction (13).

Even though they are defined in the same system, obscurus and omnia entropy are fundamentally different (e.g., at equilibrium S_*NU*_ > S_*NT*_) because the former defines the number of branches and the latter describes the number of possible combinations of branches. The branches of the obscurus are entangled in all possible combinations of branches of the omnia. If we want to understand the noise of stochastically tossed coins, we consider obscurus entropy. If we want to understand the irreversible signal, we consider omnia entropy.

Figures 2G and 2H are plots of negative entropies along molecular, *N*_*U*_, and ensemble, *N*_*T*_, reaction coordinates. Consistent with Gibbs’ chemical thermodynamics (17), the slopes of these plots describe the spontaneity of the reactions. Changes in obscurus entropy are linear with a slope of

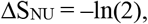

meaning that with each K → U transition, two degrees of freedom irreversibly pull the reaction forward before equilibrating at *N*_*U*_ = *N*.

In contrast, omnia entropy creates an entropic well within which a change in entropy with a chemical step, *W*_*i*_ → *W*_*f*_, from an initial i = *N*_*T*_, to final, f = *N*_*T*_+1, state is

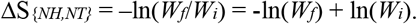

With each step, *W*_*f*_ pulls the reaction forward and *W*_*i*_ pulls the reaction backward. Substituting Eq. 1, we obtain ΔS_*NT*_ = –ln*N*_*H*_ + ln(*N*_*T*_ + 1), which simplifies the agency to: *N*_*H*_ pulls the reaction forward and, for *N*_*T*_ >> 1, *N*_*T*_ pulls the reaction backward. A net irreversible forward flux occurs when *N*_*H*_ > *N*_*T*_, and the reaction reaches equilibrium when *N*_*H*_ = *N*_*T*_. A reaction quotient, Q = *N*_*H*_/*N*_*T*_, greater than one describes the non-equilibrium entropic force that generates a net flux from H to T.

The net irreversible flux is (N_H_ – N_T_)*·k* (Fig. 1C, legend), giving a time-dependence of (*N*_*H*_ – *N*_*T*_)(t) = *N·*exp(–*k*t), or *N*_*T*_ = ½*N* – ½*N·*exp(-*k*t) (Fig. 1B, solid line). Here, omnia kinetics, *N*_*T*_(t), describe the same time course as obscura kinetics, *N*_*U*_(t) (Fig. 1B, red circles). However, only through omnia kinetics are we able to describe the relationships between the chemistry of an ensemble of molecular switches, macroscopic chemical free energies, and macroscopic force in muscle.

Here we compare omnia kinetics, *N*_*T*_(t), with the observed kinetics, *n*_*T*_(t), described in textbooks.

### Comparison to Conventional Chemistry

When we observe molecular transitions between H and T (Fig. 2B, black lines), we typically do not observe coin tosses (Fig. 2B, red circles), and so chemists assume that branched trajectories and unknown states, U, do not exist. With this assumption, chemists linearize unpredictable branched trajectories, defining reversible H to T transitions along a linear molecular reaction coordinate (Fig. 1A). Forward and reverse reaction rates are defined as *k* weighted by H to T and T to H transition probabilities, while H to H and T to T transition probabilities are ignored, resulting in molecular reaction rates of ½*k*. While it is the desire of molecular determinists that obscurus mechanisms have omnia agency – i.e., that the noise, H ↔ T, in Fig. 2B (black squares) describes the signal – this would require that we define two different entropies (obscurus and omnia) at the same time, which is not possible.

The numbers of coins, *n*_*H*_ and *n*_*T*_, observed after coins have been tossed do not have deterministic agency. Nevertheless, chemists create chemical potentials and a molecular reaction quotient, *n*_*H*_/*n*_*T*_, (a concentration gradient) (26) to account for omnia kinetics on an obscurus scale. Like other deterministic constructs (e.g., Boltzmann’s H function, the kinetic theory of gases, and molecular models of biological function) chemical activities project the agency of the omnia, *N*_*H*_/(*N*_*T*_+1), on to molecules, *n*_*H*_/*n*_*T*_. Defining obscurus and omnia entropies at the same time results in paradoxes in science that are often described as spooky (see below). However, this spookiness results from the false assumption that *n*_*T*_(t) determines *N*_*T*_(t). There is nothing spooky about the statistics of unknown branched trajectories being mathematically entangled in the statistics of all possible combinations of branched trajectories, *N*_*T*_(t), which are predictive of *n*_*T*_(t).

The reaction coordinate is linearized – not by ignoring unpredictable molecular branches but – by considering all possible branches. The obscurus arrow of time irreversibly pulls molecules into disordered branches (Figs. 2E and 2G), whereas the omnia arrow of time irreversibly pulls ensembles into all possible combinations of molecular branches (Fig. 2F) in one direction down an entropic well (Fig. 2H) toward relatively ordered states. The arrow of time does not push molecules down observed linearized molecular paths (Fig. 2A). An irreversible flux from H to T is the effect, not the cause, of the omnia reaction, *N*_*H*_(t), and is irreversibly pulled forward by an increase in entropy.

### Implications

The statistical analysis above applies to any system on any spatiotemporal scale within which unpredictable interventions randomly create branches in the trajectories of components of that system. This includes stellar systems, social systems, economic systems, political systems, ecological systems, and the evolution of life on earth. In all cases, an irreversible system trajectory is defined by an increase in omnia entropy and not by obscurus mechanisms. This has profound implications for predictive models across all disciplines.

While thermal energy randomly chooses a path at each branch point, it does not push molecules down a specific path (entropy pulls). Any other form of energy, including light energy, can do the same for branched trajectories that exist on the scale of that energy. In muscle, the coins are motor proteins that function as protein switches randomly tossed by thermal energy, k_B_T. For ten motor switches, the entropic component to the free energy of state {5,5} is k_B_T·ln(252) (Fig. 2H), which irreversibly pulls muscle into this contracted state. This is the foundation of a simple binary model that accurately describes most key aspects of muscle contraction (13).

The above analysis demonstrates that the mechanism of irreversible ensemble processes is not molecular, disproving all molecular models of irreversible ensemble processes, including the kinetic theory of gases. Briefly, for an ideal gas consisting of *N* gas molecules with 3 degrees of freedom each, the omnia entropic energy, k_B_T·ln(3^N^), is balanced against PV, or PV ≈ Nk_B_T. Analogous to the evolution of a binary distribution with time (above) or the contraction of muscle (13), the mixing or expansion of gases is not pushed by gas molecules; it is irreversibly pulled by an increase in the number of possible branches, *W*, created with mixing or expansion, solving Loschmidt’s paradox.

The transformation in scale from obscurus to omnia is energetic as well as entropic and kinetic (see Table I). In muscle, the free energy of molecular switches becomes the enthalpy of muscle, and the number of microstates of molecular switches becomes the entropy of muscle (27). The ideal gas equivalent would be that the kinetic energy of gas molecules becomes the enthalpy (heat) of the gas, and the number of microstates of gas molecules becomes the entropy of the gas.

**Table 1.** Molecular and Ensemble Scales of a Two-State Chemical Relaxation.

|  |  |  |
| --- | --- | --- |
| **Definitions** | **Molecular (Obscurus)** | **Ensemble (Omnia)** |
| States | $N_U, N_K$ | $\{N_H, N_T\}$ |
| Useful | No | Yes |
| Predicts | Disorder/Noise | Order/Signal |
| Degrees of Freedom | 2 (branched) | 1 (Linear Reaction Coordinate) |
| Entropy | $N_U \ln(2)$ | $\ln W = \ln[N!/(N_H!N_T!)]$ |
| Equilibrium | $N_U = N$ | $\{\frac{1}{2}N, \frac{1}{2}N\}$ |
| Reaction Quotient | Not Defined | $W_f/W_i = N_H/(N_T + 1)$ |
| Reaction Rate | $N_K \cdot k$ | $(W_f - W_i) \cdot k = (N_H - N_T - 1) \cdot k$ |

The non-scalable omnia and obscurus entropies create component-ensemble dualities, resembling the particle-wave duality of light, only defined on any scale. This solves the paradox of Maxwell’s demon because it precludes an obscurus demon from having omnia agency.

Similarly, it demonstrates that obscurus structural mechanisms often described in molecular biology cannot describe the omnia agency of biological function. We have shown how this solves the Gibbs paradox (19), and it appears to solve the paradox of Schrodinger’s cat. That is, if an omnia system is defined at a given instant of time, the obscurus states of its components can only be defined probabilistically at that same time.

Omnia entropy challenges the conventional view that molecular mechanisms determine irreversible ensemble mechanisms. Indeed, irreversible omnia agency (ensemble states a priori defined that pull us into the future) more closely resembles the agency of gods than molecular determinists. While some might claim that omnia entropy breaks predictive science; we contend it mends a broken science and anticipate that it will lead to significant scientific advances. A predictive entropic model of muscle contraction (13, 14) is just the beginning.

## Acknowledgments

We are grateful for the many brilliant scientists, mentors, students, and administrators who have guided and supported this work, including A.V. Hill, L. Boltzmann, J.W. Gibbs, D.D. Thomas, D.M. Warshaw, C. Cremo, K. Sanders, D.R. Jackson, A. Hooft, V. Murthy, T. Stewart, K. Cox, T. Schwenk, J. Garfield, and C. Song

